# Embodied precision: Intranasal oxytocin modulates multisensory integration

**DOI:** 10.1101/361261

**Authors:** Laura Crucianelli, Yannis Paloyelis, Lucia Ricciardi, Paul M Jenkinson, Aikaterini Fotopoulou

## Abstract

Multisensory integration processes are fundamental to our sense of self as embodied beings. Bodily illusions, such as the rubber hand illusion (RHI) and the size-weight illusion (SWI), allow us to investigate how the brain resolves conflicting multisensory evidence during perceptual inference in relation to different facets of body representation. In the RHI, synchronous tactile stimulation of a participant’s hidden hand and a visible rubber hand creates illusory bodily ownership; in the SWI, the perceived size of the body can modulate the estimated weight of external objects. According to Bayesian models, such illusions arise as an attempt to explain the causes of multisensory perception and may reflect the attenuation of somatosensory precision, which is required to resolve perceptual hypotheses about conflicting multisensory input. Recent hypotheses propose that the precision or salience of sensorimotor representations is determined by modulators of synaptic gain, like dopamine, acetylcholine and oxytocin. However, these neuromodulatory hypotheses have not been tested in the context of embodied multisensory integration. The present, double-blind, placebo-controlled, crossed-over study (N = 41 healthy volunteers) aimed to investigate the effect of intranasal oxytocin (IN-OT) on multisensory integration processes, tested by means of the RHI and the SWI. Results showed that IN-OT enhanced the subjective feeling of ownership in the RHI, only when synchronous tactile stimulation was involved. Furthermore, IN-OT increased the embodied version of the SWI (quantified as weight estimation error). These findings suggest that oxytocin might modulate processes of visuo-tactile multisensory integration by increasing the precision of top-down signals against bottom-up sensory input.

## Introduction

When we wake up in the morning, we do not question whether our body belongs to us. However, the ability to recognise our body as our own (i.e. sense of body ownership) and the consequent mental representation of our body (i.e. body image) are not axiomatic, and they both contribute to create a coherent body representation (Gallagher, 2000; Blanke, 2012). Body image involves a conscious process to identify the body as our own, and therefore, it is closely related to our sense of body ownership. It includes visual perceptions and beliefs about our own body, and it is considered to be a fundamental aspect of bodily self-consciousness (see Blanke, 2012 for a review; Dijkerman, 2015).

Experimental methods of multisensory integration allow us to investigate these distinct facets of body representation. For example, the development of experimental paradigms that allow the controlled manipulation of limb ownership in laboratory settings, such as the rubber hand illusion (RHI, Botvinick & Cohen, 1998), provide a unique tool to investigate the sense of body ownership. In this illusion, synchronous touch between a visible rubber hand and the participant’s hidden hand produces the illusion of ownership over the fake hand. This feeling of ownership towards the rubber hand arises as a solution to the unlikely conflict between sensory signals from three modalities that need to be integrated - vision, touch and proprioception (i.e. synchronous vision and touch but incongruent proprioception). People select the most plausible explanation for the causes of these different sensations (i.e. *the [rubber] hand I see being touched in synchrony with my [real] hand is most likely to be mine*) by placing greater weight to visual signals (i.e. *the hand I see*) relatively to the incompatible proprioceptive and tactile information (i.e. *where I feel my hand and the touch to be)* (Tsakiris and Haggard, 2005; Apps and Tsakiris, 2014). Recent studies have further shown what when proprioception or somatosensation are damaged or become unreliable, this inferential process continues to the point that mere vision of a rubber hand in peripersonal space is sufficient to create strong feelings of rubber hand ownership (a phenomenon termed ‘visual capture of ownership’; Martinaud et al., 2017). Taken together, RHI studies in healthy and clinical populations suggest that the sense of body ownership arises as a consequence of the attempt to find the most likely cause of multisensory signals, depending on their reliability (Apps and Tsakiris, 2014; Zeller et al., 2014).

The experience of the RHI can affect both the representation of our own body as well as the representation of the material and physical world beyond our body, e.g. the perception of objects (Haggard and Jundi, 2009). By means of a RHI paradigm, Haggard and Jundi (2009) manipulated body image representation, first explicitly, by inducing participants to embody either a larger or a smaller rubber hand than their own hand. Participants were equally able to embody the larger and small rubber hand. Second, they tested whether participants were able to judge the weight of external objects having constant size but different weight, which leads to an embodied version of the size-weight illusion (SWI, Cesari and Newell, 1999; Haggard & Jundi, 2009). Typically, in such illusions the size of an object will influence our expectations of how heavy this object is (Flanagan and Beltzner, 2000). In this embodied version of the illusion, the size of the object itself is not varied but by varying the size of the embodied hand, any changes in the perception of the object weight are considered a result of the relative size of the object in comparison to the participants’ ‘own’ hand (Haggard & Jundi, 2009). The results revealed that while there was no effect of hand size on explicit feelings of embodiment, the represented size of the participants’ hand had an impact on the perceived size and weight of the grasped object, inducing a SWI. In particular, only when participants acquired ownership of a larger hand, they perceived the grasped objects as smaller in size and therefore heavier in weight. This phenomenon can be explained by a violation of perceptual experience due to a mismatch between what we expect (e.g. the size of the object) and what we experience with our senses (e.g. the weight of the object) (Flanagan and Beltzner, 2000; Haggard and Jundi, 2009). Interestingly, the latter task can be considered as an indirect measure of body size representation, in the sense that an induced variation in the perceived body size can be measured indirectly, i.e. not by asking participants but by measuring the modulation of object weight perception. Hence, including this measure can reveal whether IN-OT acts differently at an explicit or implicit level of body representation.

These multisensory integration phenomena appear to relate to certain psychometric characteristics, such as eating disorders symptomatology and self-objectification. Indeed, people with eating disorders seems to be particularly susceptible to bodily illusions, showing multisensory integration also when tactile and visual stimulations are presented out-of-synchrony, for example (Eshkevari et al., 2012). This phenomenon persists even after otherwise successful recovery (Eshkevari et al., 2015), and might suggest the presence of a different modality of multisensory integration, which seems to be mainly visually driven. In other words, people with eating disorders tend to attribute greater weight to top-down information (e.g. the hand they see) compared to bottom-up somatosensory signals (e.g. tactile stimulation on their own hand). Recent studies suggest that (non-clinical) eating symptomatology might relate to body satisfaction following manipulation of illusory body size (Preston and Ehrsson, 2014), suggesting that this measure might influence body size representation in healthy samples. Similarly, the extent to which individuals experience their own body as an object to be evaluated based on its appearance rather than effectiveness (i.e. self-objectification), is related to eating disorders symptomatology and seems to affect body shame, anxiety and eating disorders, as well as being a potential precursor to depression and sexual dysfunction (Friedrickson and Roberts, 1997). Additionally, self-objectification accounts for the relative insensitivity of women to their own internal bodily cues, which has been reported in studies of interoception (Tiggemann and Lynch, 2001; Ainley and Tsakiris, 2013).

Computational approaches to multisensory integration (Ernst & Banks, 2002) illusions such as the RHI (Samad et al., 2015) and the SWI (e.g. Buckingham & Goodale, 2013) have shown that cross-modal conflicts are resolved in a Bayes-optimal manner, by weighting the various sensory signals with an appropriate level of confidence or precision (O’Reilly et al., 2012; Zeller et al., 2016). This concept is central to predictive coding theories of perception, which claim that the weighting of ambiguous or noisy sensory signals determines how they are used to update predictions about their sources (Friston and Stephen, 2007; Friston, 2008). Specifically, predictive coding is based on the idea that the brain interprets sensory information according to a hierarchical generative model of the world, which generates top-down representations, against which bottom-up sensory signals are tested to update beliefs about their causes (e.g., Rao & Ballard, 1999; Friston, 2005; Limanowski and Blankenburg, 2015). In this context, perception can be considered as the process of minimizing prediction errors (i.e. difference between predictions and sensory signals) by forming Bayes-optimal, top-down predictions. Crucially, the balance and reciprocal influence between bottom-up signals and top-down predictions depends on their relative uncertainty (or formally its reverse, precision) (Feldman and Friston, 2010) or their reliability (Rohe and Noppeney, 2015; Deroy et al., 2016). For example, using modelling under a predictive coding framework, Zeller and colleagues suggested that in order to resolve the uncertainty between the conflicting visual, proprioceptive and tactile information in the RHI, the brain downregulates the precision of ascending somatosensory prediction errors so that they will have less influence on top-down, predicted sensory signals about the body (Zeller et al., 2014; 2016). In support of this prediction, they found that touch-evoked EEG potentials elicited by brush-strokes during the RHI are selectively attenuated by reducing their precision (Zeller et al., 2014). These results are consistent with the idea that multisensory integration requires weighting of sensations according to their reliability, such that in the RHI the precision of somatosensory signals needs to be attenuated relative to the precision of visual signals in order to resolve conflicting perceptual hypotheses about the most likely cause of sensations in favour of the visually-perceived rubber hand (Zeller et al., 2014; 2016).

Similarly, predictive coding models of the SWI suggest that this illusion is the product of perceptual processes specialised for the detection and monitoring of *outliers* in the environment with the final goal of efficient coding of information (Buckingham & Goodale, 2013). In Bayesian terms, our weight perception is driven by our expectations of how heavy something should be. These priors are weighted against bottom up signals and adjusted by the presence of lifting errors. When lifting an object for the first time, participants will apply either excessive or insufficient force; however, in subsequent trials, it is likely that previous experience will reduce the amount or size of such errors (Buckingham & Goodale, 2013).

These Bayesian accounts of multisensory integration seem to suggest that the brain is able to represent and use estimates of uncertainty (more formally *precision*, the reverse of uncertainty; Friston, 2010; Knill & Pouget, 2004) in order to achieve an optimal coding of information, even if the underlying neural coding principles remain debated (e.g. Ma et al., 2006 vs. Fiser et al., 2010). According to one view, precision is thought to be mediated by the gain or excitability of (superficial pyramidal) cells encoding prediction errors (Feldman and Friston, 2010; Bastos et al., 2012; Shipp et al., 2013). Thus, optimizing precision corresponds to neuromodulatory gain control of neuronal populations reporting prediction error (Feldman & Friston, 2010). On this hypothesis, modulators of synaptic gain (like dopamine, acetylcholine and norepinephrine, as well as neuropeptides such as oxytocin, Quattrocki & Friston, 2014) therefore might play a role in determining the precision or salience of sensorimotor representations encoded by the activity of the synapses they modulate (Fiorillo, Tobler & Schultz, 2003; Yu and Dayan, 2005; Friston et al., 2012). However, to our knowledge these hypotheses have not been tested in multisensory integration research. The present study specifically aimed to study the role of the neuropeptide oxytocin on multisensory integration and particularly body ownership and body image representation.

Oxytocin is a neuropeptide consisting of nine amino acids, mostly synthesised in the hypothalamus. This neuropeptide acts peripherally as a hormone, such as when it is released during labor and breastfeeding to stimulate uterine contractions and milk ejection, respectively (Burbach, Young & Russell, 2006). Oxytocin also acts centrally as a neuromodulator (Stoop, 2012; Grinevich et al., 2016; Uvnäs-Moberg et al., 2015). It has been proposed that oxytocin might modulate the relation between bottom-up sensory information and top-down attentional biases by enhancing the salience of socially relevant information and attenuating the saliency of non-social stimuli (Gordon et al., 2013; Quattrocki & Friston, 2014). Oxytocin might interact with the dopaminergic system in order to increase the attention orientation towards social cues (Shamay-Tsoory & Abu-Akel, 2016). The use of nasal sprays has provided a safe and noninvasive route for the administration of oxytocin to target the human brain (MacDonald, Dadds, Brennan, Williams, Levy et al., 2011; Born et al., 2002; Paloyelis et al., 2016). An increasing number of studies have proposed that intranasal oxytocin can modulate social cognition, affiliation and brain function in humans (De Dreu et al., 2010; MacDonald & MacDonald, 2010; Colonnello and Heinrichs, 2016; Uvnäs-Moberg et al., 2015; Rocchetti et al., 2014).

Here, in a parsimonious psychophysical setting, we examined whether (1) oxytocin affects body ownership during a classic RHI; and (2) oxytocin modulates body image representation, by means of an enhanced version of the illusion combining the RHI with hands of different sizes and a SWI (as in Haggard and Jundi, 2009). In both instances, individuals had to weigh conflicting sensations deriving from the self (somatosensory signals) or rubber hands of different sizes (visual signals). Specifically, based on theoretical proposals about the role of oxytocin on enhancing somatosensory attenuation (Quattrocki & Friston, 2014), we hypothesised that oxytocin would enhance the embodiment of the rubber hand in the standard and in the enhanced RHI paradigm, by attenuating the precision of somatosensory signals relative to the precision of visual signals. In the standard RHI paradigm (1), we predicted that IN-OT would modulate multisensory integration and corresponding subjective (i.e. self-report measure) and behavioural (i.e. the degree to which participants erroneously perceived changes in the position of their own, unseen hand towards the rubber hand, the so-called proprioceptive drift) measures of body ownership over the RHI. We expected to observe these effects in both mere visual capture conditions and during synchronous tactile stimulation of the rubber hand and participants’ own hand. In the SWI combined with the RHI (2), we predicted that IN-OT would have an effect on body image representation by increasing the weight estimation error following embodiment of the larger hand; in contrast, no effect on weight estimation was expected following embodiment of the normal size hand.

In sum, we conducted a double blind, placebo-controlled, crossover study to investigate the effect of a single dose of IN-OT compared to placebo on multisensory integration during a standard and an enhanced version of the rubber hand illusion paradigm where we manipulated the synchronicity of touch (synchronous vs. asynchronous) and the size of the rubber hand (normal vs. large). We recorded subjective and behavioural measures of body ownership in both mere visual capture conditions and during synchronous tactile stimulation of the rubber hand and participants’ own hand. We measured body image representation by means of weight estimation errors during the SWI. Measures of eating disorders symptomatology (Eating Disorders Examination Questionnaire, EDE-Q, Fairburn & Beglin, 1994) and self-objectification (Self-Objectification Questionnaire, Fredrickson et al., 1998) were also included to control for their potential role in standard processes of multisensory integration.

## Materials and Methods

### Participants

Forty-one heterosexual females were recruited through the University College London research participation system. They were aged between 18 and 40 years (M = 25.13, SD = 4.21). Participants were not taking any medication (including the contraceptive pill) and were recruited in the follicular phase of their menstrual cycle (between the 5^th^ and 14^th^ day) to control for hormonal levels (Salonia et al., 2005). Exclusion criteria included being left handed, pregnant or currently breastfeeding (see MacDonald et al., 2011), a history of any medical, neurological or psychiatric illness, BMI out of the range 18.5 – 24.9 (M = 21.38; SD = 2.64), use of any illegal drugs within the last six months, and consumption of more than five cigarettes per day. Participants were asked to refrain from consuming any alcohol the day before testing and any alcohol or coffee on the day of testing. All participants provided informed consent to take part and received a compensation of £40 for travelling expenses and time. Ethical approval was obtained by University College London and the study was carried out in accordance with the provisions of the Helsinki Declaration of 1975, as revised in 2008. One participant was excluded because she discontinued the study (i.e. she took part in only one of two testing sessions); four participants were later excluded from the analysis as they were found to be extreme outliers in their embodiment scores in the placebo condition (embodiment score ± 3 SD from group mean). The final sample comprised 36 participants (M age = 25.03, SD = 3.96).

### Experimental design

The study employed a double-blind AB/BA (oxytocin-placebo/placebo-oxytocin) cross-over design, with compound (oxytocin vs. placebo) as the within-subjects factor. Each subject participated in two identical sessions, each lasting between 1.5 and 2 hours and conducted between 1 and 3 days apart; this was to ensure that they were tested in the same phase of the menstrual cycle. The testing sessions always took place between 09.00 and 12.00. In one session, participants were asked to self-administer 40 IU of oxytocin and in the other session placebo (see Figure 1) in a counter-balanced and double-blinded manner. Participants were randomly allocated to the nasal spray sequence (AB/BA); eighteen participants received placebo on the first visit and intranasal oxytocin on the second visit, while another 18 received the reverse order (intranasal oxytocin on the first visit and placebo on the second visit). The order of administration was included as a covariate in all analyses. The rubber hand illusion and size weight illusion (see below for details of procedure) were administered 30 minutes after nasal spray administration (see MacDonald et al., 2011; Paloyelis, Doyle, Zelaya, Maltezos, Williams et al., 2016 for optimal temporal window). The asynchronous condition of the rubber hand illusion was run only once to establish the presence of the illusion, while the synchronous condition was repeated twice; once with a “normal” size hand and once with a “larger” hand (see below for more details) to investigate the effect of hand size on the embodiment process. The order of the three conditions (normal hand/synchronous, normal hand/asynchronous, larger hand/synchronous) was pseudorandomised between participants (i.e. such that the larger hand was never administrated as the first condition so to always obtain the first visual capture measure with the normal size hand) but it was kept constant within participants between the two testing session.

**Figure 1.**
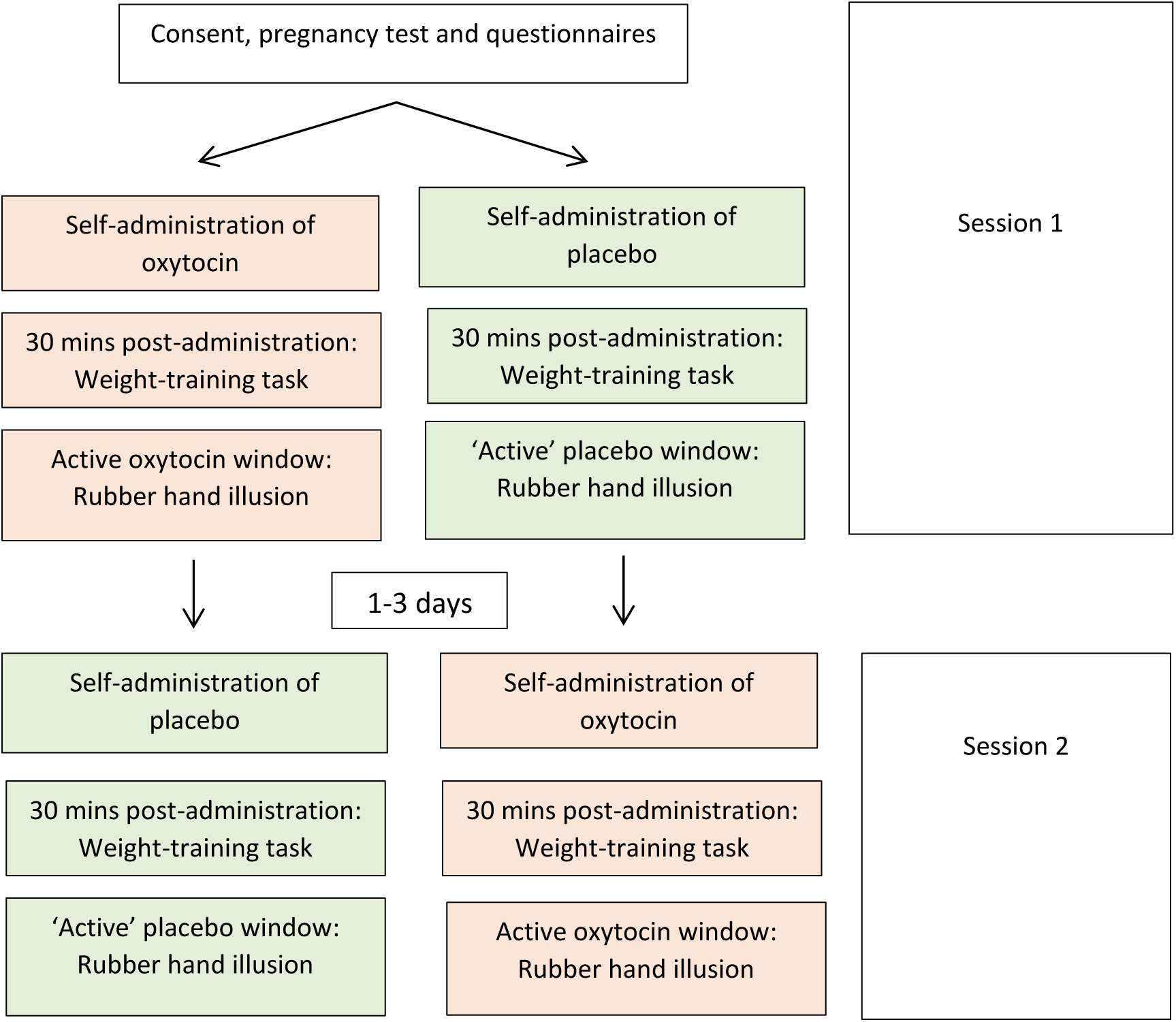
Study design and flowchart.

#### Oxytocin and placebo spray

Participants received 40IU of oxytocin (Syntocinon, Novartis Pharmaceutical, Basel, Switzerland) or placebo (containing the same ingredients as Syntocinon except the active ingredient) by means of a nasal spray. Two practice bottles containing water were used for the participants to familiarise themselves with the procedure; one for the experimenter to demonstrate the correct technique and one for the participant to practice. Experimental instruction about the nasal spray administration procedure, the position of the head and of the nasal spray inside the nose, and breathing technique were given to the participants. Before the beginning of the self-administration procedure, all the participants were asked to blow their nose. Participants self-administered a puff containing 4IU every 30 seconds alternating between nostrils (five for each nostril) for a total of ten puffs. Half of the sample started the administration on the right nostril, and half on the left nostril. Participants were specifically instructed to not blow their nose during the administration procedure. At the end of the last puff of nasal spray, participants were given three minutes of resting time in which they were instructed to relax. The self-administration procedure took approximately nine minutes, including three minutes of rest at the end (see also Paloyelis et al., 2016).

### Rubber hand illusion

#### Procedure

The rubber hand illusion was performed following the procedure fully described in previous studies (Crucianelli et al., 2013; Crucianelli et al., 2017). Two adjacent stroking areas, each measuring 9cm long x 4cm wide were identified and marked with a washable marker on the hairy skin of participants’ left forearm (wrist crease to elbow, McGlone, Olausson, Boyle, Jones-Gotman, Dancer et al., 2012). Tactile stimulation (i.e. stroking) was alternated between these two areas to minimise habituation, and congruent stroking area changes were applied to the rubber hand in all instances (Crucianelli et al., 2013). In each condition, the experimenter placed the participant’s left hand (palm facing down; fingers pointing forwards) at a fixed point inside a wooden box (34 cm × 65 cm × 44 cm).

A pre-stroking estimate of finger position was then obtained (for the measurement of proprioceptive drift) using a tailor’s tape-measure placed on top of the box lid. Subsequently, the rubber arm was positioned in the right half of the box (participant-centred reference frame), in front of the participant’s body midline, and in the same direction as the participant’s actual left arm. The distance between the participants’ left arm and the visible arm (on the sagittal plane) was approximately 28 cm. A wooden lid prevented visual feedback of the participant’s own arm. The participant also wore a black cape to occlude vision of the proximal end of the rubber arm and participant’s left arm. The rubber hands were custom made by an artist with expertise in body-part modelling, by means of hand casts obtained from two volunteers, one with a BMI within the healthy weight range (healthy BMI = 18.5-24.9), and one having a BMI in the obese range (obese BMI = 30-39.9). Each cast was subsequently hand painted to create a life-like model hand (Figure 2).

**Figure 2.**
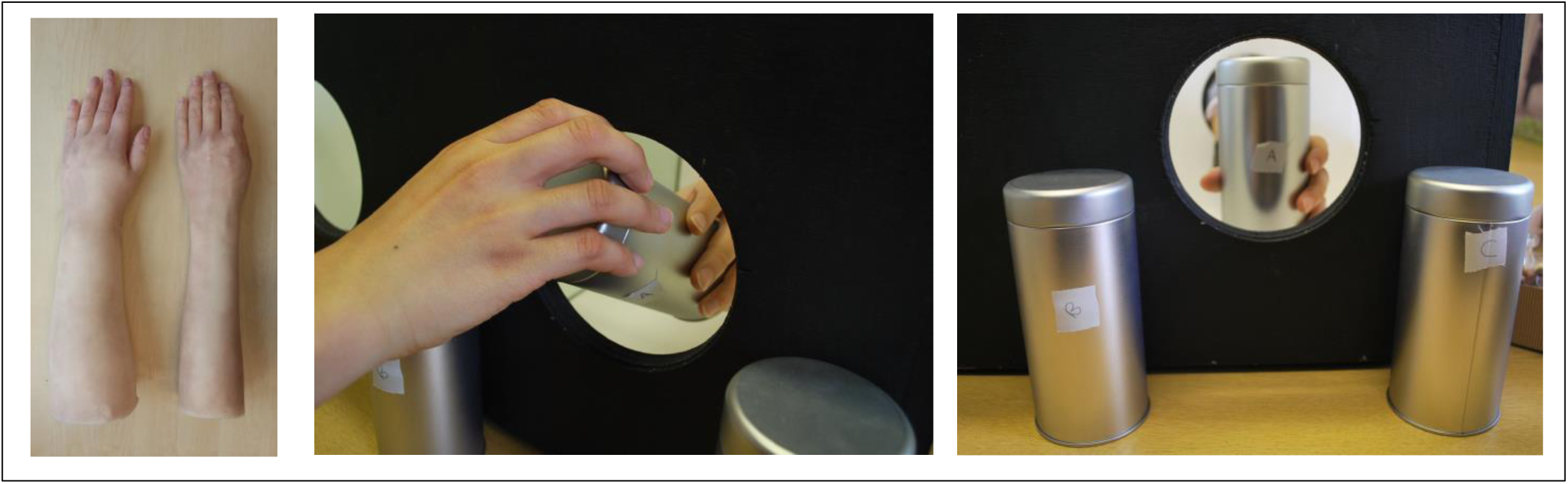
The size-weight illusion procedure. Firstly, participants were completing a RHI procedure where they were asked to look at a regular size rubber hand (on the right) or a larger size rubber hand (on the left). Participants were asked to estimate the weights of three cans, having identical size but different weight (i.e. 125g, 150g, and 175g). The experiment was passing the can through the hole in the box. Participants were instructed to lift the can, without shaking it, and to give the best estimation about their weight. No feedback was provided to the participants.

Participants were then asked to look at the rubber hand continuously for 15 seconds, before completing the pre embodiment questionnaire (i.e. *visual capture effect*). Tactile stimulation was then applied for one minute using two, identical, cosmetic make-up brushes (Natural hair Blush Brush, N°7, The Boots Company) using the speed of 3 cm/s (Crucianelli et al., 2013). In the synchronous conditions, the participant’s left forearm and the rubber forearm were stroked such that visual and tactile feedback were congruent, whereas in the asynchronous conditions, visual and tactile stimulation were temporally incongruent (i.e. offset by 2 seconds). After the stimulation period, the felt and actual location of the participant’s left index finger was again measured following the pre-induction procedure. After the tactile stimulation period, participants completed the post-stroking *embodiment questionnaire.* Prior to commencing the next condition, they were given a 60s rest period, during which they were instructed to freely move their left hand.

#### Outcome measures

After positioning the hand inside the box, participants were asked to close their eyes and to indicate on the ruler with their right hand the position where they felt that their own left index finger was inside the box. The experimenter then measured and recorded the actual position of the participant’s left index finger. After the stimulation period, the felt and actual location of the participant’s left index finger was again measured following the pre-induction procedure. The difference between the pre and post error in location represents a measure of *proprioceptive drift*.

An *embodiment questionnaire* (Longo et al., 2008) was used to capture the subjective experience of the illusion (12 statements rated on a 7-point Likert-type scale; −3 = strongly disagree, +3 = strongly agree). In each condition, the questionnaire was administered pre- (i.e., embodiment due to the visual capture effect) and post-stroking and we calculated their difference to obtain a measure of subjective embodiment due to visuo-tactile integration (i.e. *change in embodiment*, as in Crucianelli et al., 2013; 2017). This questionnaire consists of three sub-components: *felt ownership*, that is related to the feeling that the rubber hand is part of one’s body; *felt location* of own hand, that is related to the feeling that the rubber hand and one’s own hand are in the same place; *affect*, that includes items related to the experience being interesting. We examined this difference between pre- and post-stroking (*change in embodiment*) for each of the statements separately, as well as for an overall “embodiment of rubber hand” (Longo et al., 2008) score, that was obtained by averaging the scores of the two subcomponents specifically related to embodiment, namely ownership and felt location that did not relate to affect. The affect sub-component also included a measurement of the perceived pleasantness of the tactile stimulation quantified by means of a visual analogue scale ranging from 0 (not at all pleasant) to 100 (extremely pleasant).

### Size-weight illusion

#### Procedure

The size-weight illusion task was based on the procedures described by Haggard and Jundi (2009). Before the main experiment (see timeline in Figure 1), participants received training in weight estimation. They were asked to lift two opaque cylinders (i.e. metal tea caddies/containers, Figure 2) that contained wooden beads. Each cylinder was the same size (height = 15.5; diameter = ~7.5cm) but varied in weight (either 100g or 200g). These same cylinders were later used in the actual experimental task. Participants were asked to estimate the weight of the cylinders ten times by lifting them with their left hand and putting them back down; five times they were given the cylinder weighting 100g, and five times they were given the cylinder weighting 200g, in a random order. During this training stage participants were given feedback about their performance (i.e. if they were right or wrong) and were informed about the actual weight of the cylinder after each trial. By the end of the training session, all participants successfully learnt to distinguish between cylinders weighting 100g and 200 g (i.e. the cut off was 8 out of 10 correct trials in a row). This training allowed participants the opportunity to familiarise themselves with the experimental stimuli and ensured that all participants started the main experiment with the same weight reference points (see also Haggard and Jundi, 2009).

Subsequently the baseline weight-estimation took place. The weight estimation task consisted on lifting three cans of the same shape and size (described above), but different weight (125g; 150g; 175g, presented in a random order across conditions and participants). The experimenter placed the can between the participants’ left index finger and thumb. Participants were asked to gently lift the can without shaking it, to put it down, and to provide verbally the best estimation about its weight (to the nearest gram). Instructions included the information that the weight could be anything between 100 and 200g but not exactly 100 or 200 g; no feedback about the estimation was given at this stage. During the session, participants completed the weight-estimation task twice more: following the rubber hand illusion with the normal size hand, and following the rubber hand illusion with the larger size hand. The difference between the estimated and actual weight was the measure of the SWI (i.e. weight estimation error), which was then compared between the conditions following embodiment of the normal size or larger hand in order to investigate the relationship between expectations driven by the *illusory* body size perception and the estimated weight of the objects.

#### Self-Objectification Questionnaire (SOQ)

The Self-Objectification Questionnaire (Fredrickson et al., 1998) measures the extent to which individuals see their bodies in observable, appearance-based (i.e. objectified) terms, versus non-observable competence-based terms. The questionnaire consists of ten body attributes (e.g. attractiveness, strength, health, etc.) which participants are required to rank by how important each is to their own physical self-concept, from 0 (for least impact) to 9 (greatest impact). Self-objectification scores are calculated by subtracting the summed ranks given to the 5 competence-based attributes (e.g. health, energy) from the summed ranks of the 5 appearance-based attributes (e.g. physical attractiveness, body measurements). Scores range from 25 to 225, with higher scores indicating greater emphasis on appearance, which is interpreted as greater self-objectification. The SOQ has good test-retest reliability (r =.92, cited in Miner-Rubino, Twenge and Fredrickson, 2002).

#### Eating Disorders Examination Questionnaire (EDE-Q)

The Eating Disorder Examination Questionnaire 6.0 (EDE-Q; Fairburn & Beglin, 1994), which has good consistency and reliability (global score α = .90; Peterson et al., 2007), was used to measure eating disorders symptomatology. The questionnaire consists of 28-items rated on a seven-point Likert scale (ranging from *No days/Not at all* to *Every day/Markedly*), and six items asking about frequency of behaviour. The questionnaire can be divided into four subscales (*dietary restraint*, e.g. Have you been deliberately trying to limit the amount of food you eat to influence your shape or weight?; *eating concern*, e.g. Over the past 28 days, how concerned have you been about other people seeing you eat?; *weight concern*, e.g. Has your weight influenced how you think about yourself as a person?; *shape concern*, e.g. Have you had a definite desire to have a totally flat stomach?), or a single global measure. To obtain an overall or “global” score, the four subscales scores are summed and the resulting total divided by the number of subscales (i.e. four). The clinical cut-off is of 2.8 on the global score (Mond et al., 2008).

### General Procedure

The experiment was run by two female experimenters. Upon arrival, in the first session only, participants were asked to provide a urine sample for a pregnancy test (Pregnancy test device, SureScreen Diagnostics), which was carried out to exclude the possibility of any ongoing, unknown pregnancy that might be adversely affected by the administration of oxytocin. After confirmation of the negative result of the pregnancy test, participants completed the EDE-Q and the SOQ (presented in a random order), and their arm was prepared for later administration of the rubber hand illusion (see Rubber Hand Illusion section above). Subsequently, participants self-administered, under both experimenters’ supervision, either intranasal oxytocin or the placebo. The order of the treatment was counterbalanced across participants, and both experimenters and participants were blind to the treatment order (as detailed in *Oxytocin and placebo spray* section above).

During the 30 minutes time post-spray administration (see MacDonald et al. 2011; Paloyelis, Doyle, Zelaya, Maltezos, Williams et al., 2016 for optimal temporal window) no social contact between the participant and the experimenters took place beyond necessary experimental instructions. Participants were asked to refrain from checking their phones or doing any personal reading. During this waiting time of oxytocin activation, participants were familiarised with the weight estimation task, before completing the three weight estimations baselines (described in Size-Weight Illusion section above). In the remaining time, participants were offered the opportunity to complete a Sudoku or Word Search. At the beginning of the active oxytocin window, participants completed the rubber hand illusion and the size weight illusion for the following 40 minutes. Participants were fully debriefed and reimbursed £40 for their time at the end of the second study visit.

### Statistical analysis

The present study aimed to explore whether IN-OT versus placebo would affect (1) the subjective feeling of ownership towards the rubber hand, as measured using an embodiment questionnaire, (2) proprioceptive judgements regarding the location of the positon of the real hand relative to the rubber hand (proprioceptive drift), and (3) the representation of body size, as measured by means of a SWI, quantified as weight-estimation error after the embodiment of rubber hands of different sizes.

Preliminary correlational analyses were conducted to investigate the relationships between the psychometric measures, namely Self-Objectification Questionnaire (SOQ) and Eating Disorders Examination Questionnaire (EDE-Q), with the subjective measure of the RHI, the proprioceptive drift and the weight-estimation error. These preliminary analyses informed the inclusion/exclusion of these psychometric measures as co-variates in subsequent analyses.

The main analyses were conducted by means of linear mixed model (LMM) analyses that allow the use of both fixed and random effects in the same analysis. Specifically, the effect of IN-OT versus placebo on the rubber hand illusion was analysed by means of two separate LMM analyses. One analysis was run to test the effect of IN-OT versus placebo on the *occurrence* of the illusion (i.e. comparison between synchronous and asynchronous touch condition, and the interaction between synchronicity and compound, on measures of embodiment). The other LMM analysis was run to test the effect of IN-OT versus placebo on the *hand size manipulation* (i.e. comparison between normal and larger hand in synchronous conditions only, and the interaction between hand size and compound, on measures of embodiment). Finally, the latter analysis was repeated with the weight estimation as the dependent variable, in order to investigate the effect of IN-OT versus placebo on the occurrence of the size weight illusion.

In all these analyses, order of compound administration (oxytocin-placebo or placebo-oxytocin), was included as a covariate. The baseline weight-estimation was included in all the analyses of the size-weight illusion effect as a covariate. All data were analysed using SPSS, version 23.

## Results

### Preliminary analyses

Preliminary correlational analyses showed no significant relationships between self-objectification (SOQ) or eating disorders symptomology (EDE-Q) with the subjective measure of the RHI (embodiment questionnaire), proprioceptive drift, or the weight estimation error in the SWI. Results are reported on Table 1. Given the lack of significant relationships between the psychometric measures and the main measures of interest, SOQ and EDE-Q were not taken into account in subsequent analyses.

**Table 1.** Pearson’s correlational analyses between Self-Objectification Questionnaire (SOQ) or Eating-Disorders Examination Questionnaire (EDE-Q) and measures of embodiment. The values reported are the correlational coefficients *r* in the first raw and the *p* values in the second raw.

|  |  | SOQ | EDE-Q |
| --- | --- | --- | --- |
| ***Classic rubber hand illusion – Subjective change in embodiment*** |  |  |  |
| IN-OT synchronous touch | $r$ | -.17 | -.19 |
| | $p$ | .29 | .24 |
| IN-OT asynchronous touch | $r$ | .02 | -.00 |
| | $p$ | .89 | .98 |
| PL synchronous touch | $r$ | -.06 | -.28 |
| | $p$ | .71 | .08 |
| PL asynchronous touch | $r$ | .02 | .17 |
| | $p$ | .90 | .29 |
| ***Classic rubber hand illusion – Proprioceptive drift*** |  |  |  |
| IN-OT synchronous touch | $r$ | .04 | .01 |
| | $p$ | .80 | .94 |
| IN-OT asynchronous touch | $r$ | -.03 | -.06 |
| | $p$ | .86 | .71 |
| PL synchronous touch | $r$ | .06 | -.15 |
| | $p$ | .74 | .36 |
| PL asynchronous touch | $r$ | .10 | -.13 |
| | $p$ | .55 | .45 |
| ***Size Weight Illusion – Error in weight estimation*** |  |  |  |
| IN-OT regular hand | $r$ | .02 | -.07 |
| | $p$ | .88 | .67 |
| PL regular hand | $r$ | -.01 | .15 |
| | $p$ | .97 | .34 |
| IN-OT larger hand | $r$ | .11 | -.08 |
| | $p$ | .52 | .64 |
| PL larger hand | $r$ | .04 | .25 |
| | $p$ | .80 | .12 |

### IN-OT Effects on Body Ownership: Synchronous vs. Asynchronous Stimulation Change in Subjective Embodiment as Measured by Self-Report

In order to verify whether IN-OT affects the subjective reporting of *body ownership* during the RHI, a linear mixed model analysis was run to explore the effect of IN-OT (versus placebo) and synchronous (versus asynchronous) stimulation on the change in embodiment questionnaire scores. The order of administration was considered in the analysis as a covariate. This analysis revealed a main effect of synchronicity (*F* (1, 35) = 8.8; *p* = 0.005; *r* = 0.82, see Figure 3), with synchronous touch (M= 0.75, SE = 0.16) leading to greater embodiment compared to asynchronous touch (M= −0.01; SE = 0.18). This result confirmed the occurrence of the illusion from a subjective point of view. Nasal spray did not have a significant main effect on the change in embodiment (*F* (1, 35) = 0.48; *p* = 0.49). However, there was a significant interaction between synchronicity and nasal spray (*F* (1, 35) = 4.87; *p* = 0.034; *r* = 0.64). Bonferroni corrected post-hoc analyses (p = 0.025) revealed that IN-OT (compared to placebo) increased embodiment of the rubber hand in the synchronous (*t* (35) = 12.50; *p* = 0.001; *r* = 0.90) but not in the asynchronous condition (*t* (35) = 0.05; *p* = 0.82; *r* = 0.01).

**Figure 3.**
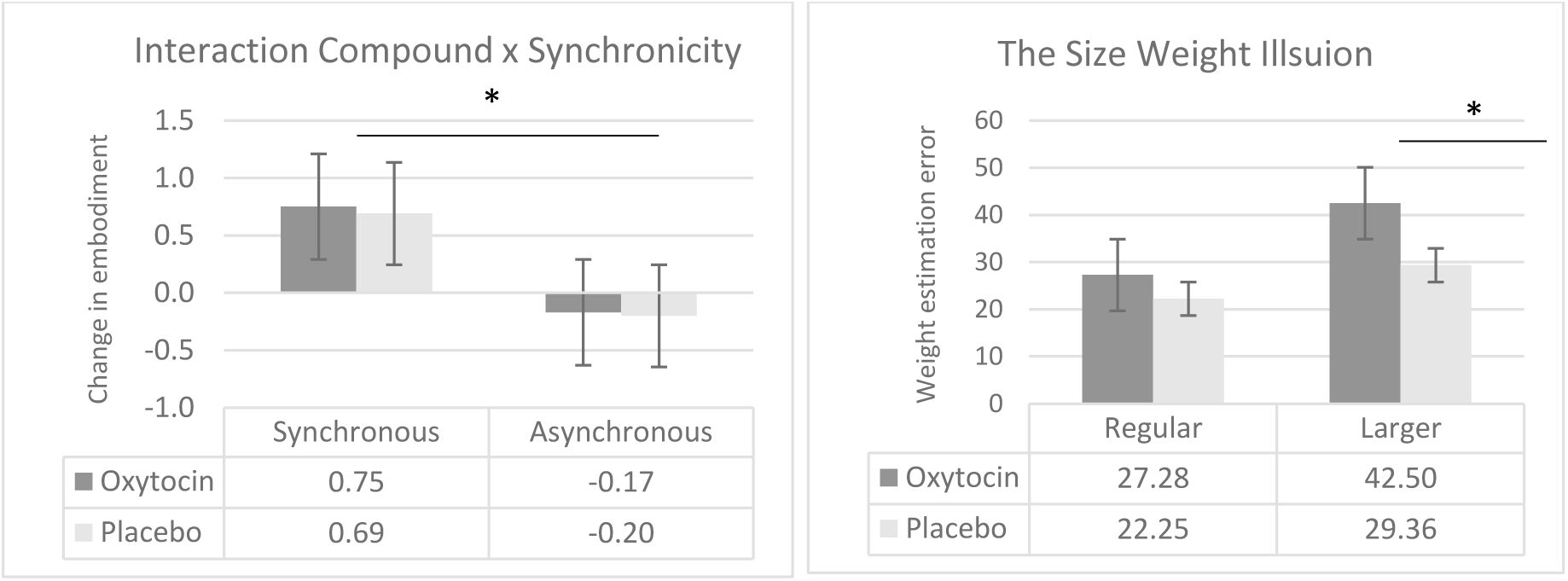
(a) Raw data mean embodiment change scores for the synchronous and asynchronous stroking conditions, after administration of oxytocin or placebo. (b) Raw data mean of weight estimation error, following embodiment of the regular and larger size rubber hand, after administration of oxytocin or placebo. Error bars denote standard error. ^∗^ indicated a *p* < 0.05

### Embodiment Changes as Measured by Proprioceptive Drift

In order to verify whether IN-OT affects the perceived location of the hand in the RHI, a linear mixed model analysis was run to explore the effect of IN-OT (versus placebo) and synchronous (versus asynchronous) stimulation on the proprioceptive drift. The order of administration was considered in the analysis as a covariate. This analysis revealed no significant main effect of synchronicity (*F* (1, 35) = 0.64; *p* = 0.43; *r* = 0.12), IN-OT (*F* (1, 35) = 0.29; *p* = 0.59; *r* = 0.04), nor a significant interaction between synchronicity and nasal spray (*F* (1, 35) = 0.51; *p* = 0.48; *r* = 0.08). Given the lack of findings regarding proprioceptive drift on our main RHI induction, we did not conduct further analyses on proprioceptive drift.

### IN-OT Effects on Embodiment of Hands of different sizes as Measured by Self-Report

In order to verify whether IN-OT affects the embodiment of hands of different sizes during the RHI, a linear mixed model analysis was run to explore the effect of IN-OT (versus placebo) and watching a larger versus regular size hand on the subjective change in embodiment. Order of nasal spray administration was considered as a covariate. This analysis showed a main effect of nasal spray (*F* (1, 35) = 10.51; *p* = 0.003; *r* = 0.87), with oxytocin leading to a greater embodiment (embodiment oxytocin, M = 0.81, SE = 0.13) compared to placebo (embodiment placebo, M = 0.74, SE = 0.14). This result is in line with the one reported above, that is IN-OT seems to enhance the embodiment of the rubber hand more than placebo regardless of the size of the hand, when touch is synchronous (i.e. no asynchronous condition was conducted with the larger hand). Indeed, hand size did not have a significant main effect on the change in embodiment (*F* (1, 35) = 3.43, *p* = 0.07; *r* = 0.25). The interaction between nasal spray and hand size was non-significant (*F* (1, 35) = 0.09; *p* = 0.77; *r* = 0.02). In sum, 1) we found that IN-OT compared to placebo enhanced the embodiment of the rubber hand, 2) we did not find an effect of hand size on embodiment of the rubber hand, and 2) we did not find that IN-OT enhances the RHI depending on rubber hand size.

### IN-OT Effects on Change in Body Size Representation: the Size Weight Illusion

Finally, a LMM analysis was run to explore the effect of nasal spray and hand size (independent variables) on the weight estimation task (dependent variable), quantified as weight estimation error. The order of administration was considered in the analysis as a covariate, together with the baseline weight estimation. This analysis showed that neither hand size (*F* (1, 35) = 0.04; *p* = 0.95; *r* = 0.01) nor nasal spray (*F* (1, 35) = 1.89; *p* = 0.18; *r* = 0.30) had a significant main effect on the weight estimation task. However, there was a significant interaction between hand size and nasal spray (*F* (1, 35) = 7.2; *p* = 0.01; *r* = 0.77, see Figure 3). Bonferroni-corrected post-hoc analyses (p = 0.025) showed no effect of oxytocin compared with placebo in weight estimation error for the regular hand (*t* (35) = −0.67; *p* = 0.51; *r* = 0.19). By contrast, oxytocin increases the weight estimation error after embodiment of the larger hand, compared with placebo (*F* (35) = −1.87; *p* = 0.007; *r* = 0.30). These results indicate that IN-OT enhances the size weight illusion in comparison to placebo when a larger hand is used, but not when a regular hand is used.

## Discussion

This double blind, placebo-controlled, crossover study aimed to explore the effect of intranasal oxytocin, in comparison to placebo, on multisensory integration in relation to the sense of body ownership (i.e. rubber hand illusion) and body image (i.e. size-weight illusion). Specifically, we investigated whether IN-OT versus placebo would affect (1) the subjective feeling of body ownership, as measured by means of the embodiment questionnaire; (2) the capture of proprioception by touch as measured by means of proprioceptive drift; and (3) the representation of body size, as measured by means of weight estimation error after embodying rubber hands of different sizes (i.e. size weight illusion).

The results showed that IN-OT enhances the subjective feeling of ownership towards a rubber hand to a greater extent compared to placebo. This effect is independent of the size of the seen hand. In fact, participants embodied the large hand to the same extent as the regular size hand, and IN-OT increases the feeling of embodiment regardless of the hand’s size. In contrast, IN-OT did not affect the proprioceptive capture of touch differently compared to placebo. Finally, this study showed that IN-OT enhanced the size-weight illusion following embodiment of a large hand only, in the sense that after administration of IN-OT participants tend to overestimate the weight of the cans to a greater extent compared to the placebo control condition. Such effect was not present following embodiment of the regular size hand.

Our findings confirmed our *first hypothesis* that embodiment of the rubber hand would be enhanced following administration of IN-OT only. Specifically, based on theoretical proposals about the role of oxytocin on enhancing somatosensory attenuation (Quattrocki & Friston, 2014), we hypothesised that oxytocin would enhance the RHI effects, by attenuating the precision of somatosensory signals relative to the precision of visual signals. Our results showed that IN-OT modulated multisensory integration and corresponding subjective measure of body ownership (i.e. embodiment questionnaire) over the RHI only during synchronous tactile stimulation of the rubber hand and participants’ own hand. No effect of IN-OT was observed in the behavioural (i.e. proprioceptive drift) measure of body ownership over the RHI, nor in the mere visual capture conditions. These findings seem to suggest that IN-OT affects the multisensory integration processes only when tactile stimulation is involved, but not in the condition when only visual and proprioceptive signals need to be integrated. In predictive coding terms, this might suggest that oxytocin, as a neuromodulator, plays a role in weighting (i.e. increasing precision) conflicting sensations deriving from the self (somatosensory signals) or the rubber hand (visual signals). This process reflects in subjective feelings of body ownership towards the rubber hand, but not in an update in the perceived location of the hand (see Rohde, Di Luca & Ernst, 2011; Abdulkarim & Ehrsson, 2016 for experimental evidence on the dissociation between these two measures of the RHI).

To the authors’ knowledge, only one recent study attempted to explore the relation between oxytocin and the sense of body ownership. Ide and Wada (2017) showed that the subjective but not objective experience of the RHI is associated with oxytocin, in the sense that participants with higher level of salivary oxytocin at baseline showed an increased experience of the RHI. The authors speculated that oxytocin might enhance the salience of tactile stimulation by modulating the activity of the insular cortex, and the higher susceptibility to the experience of the RHI would be a consequence of an increase in resources (i.e. attention) placed on touch (Ide and Wada, 2017). However, the instability of single measurements of oxytocin in peripheral fluids and the correlational nature of these findings do not allow firm conclusion to be made.

The present data also partially support our *second hypothesis* that hand size was not expected to have an effect on the rubber hand illusion *per se* (e.g. Farmer, Tajadura-Jimez & Tsakiris, 2012; Haggard and Jundi, 2009) but on the weight estimation error, here included as an indirect measure of body image (Haggard and Jundi, 2009). Our findings are in line with previous studies showing that the rubber hand illusion is object-dependent, in the sense that it occurs as long as the object could be a body part (Tsakiris et al 2010). In addition, the physical characteristics of the hand, such as skin tone (Farmer et al., 2012) do not seem to affect the occurrence of the illusion. Similarly, our findings showed that the size of the hand did not block or reduce the embodiment process towards the rubber hand. In other words, the effect of IN-OT seems to be over and above physical characteristics of the hand, and it is rather dependent on the synchronicity between seen and felt touch. Furthermore, we predicted that IN-OT would modulate multisensory integration and corresponding behavioural measure as quantified by judgments of the weight of external objects having constant size but different weight following embodiment of the larger hand only (as in Haggard and Jundi, 2009). In other words, we expected IN-OT to increase the error in weight estimation following the embodiment of a large hand only; in contrast, no effect on IN-OT on weight estimation would be observed following the embodiment of the regular size hand. Our results confirmed this hypothesis. Although participants showed a greater estimation error following the embodiment of the larger hand as compared to the regular size hand, this effect did not reach significance (see Haggard and Jundi, 2009). However, IN-OT seems to promote the occurrence of the size weight illusion compared to placebo, in the sense that it increases the weight estimation error only after embodiment of the larger hand. These findings might suggest that oxytocin allows more flexibility in terms of body shape and size. We could speculate that this process might be related to the role that oxytocin plays in pregnancy, a moment of important bodily and emotional changes in women. Additionally, the fact that IN-OT influences explicit and implicit measures of the malleability of body representation is in line with previous findings showing that oxytocin sharpens the recognition of basic emotions as well as hidden/implicit emotional facial expressions (Leknes et al., 2013; Leppanen et al., 2017 for a meta-analysis). This explanation supports the idea that oxytocin might play an important evolutionary role, by promoting social affiliation and bonding and by downregulating pain perception associated with reproduction (Insel, 1992; Paloyelis et al., 2015).

In conclusion, our findings are the first to suggest that intranasal oxytocin modulates processes of multisensory integration in relation to both body ownership and body image. We speculate that this process might follow basic principles of predictive coding. In the context of multisensory integration, oxytocin seems to mediate the weighting (or attention) given to descending prior representation and ascending sensory inputs (predictions errors). In other words, oxytocin might mediate the neurobiological mechanism promoting and sharpening the precision of sensory inputs in order to determine the weighting of top-down cognitive representations against bottom-up sensory information, ultimately facilitating multisensory integration. To our knowledge, this is the first study to have investigated whether the oxytocinergic system might modulate process of multisensory integration via embodied synchronicity mechanisms, such as synchronous tactile stimulation on the body. These findings are particularly important not only because the sense of body ownership is a fundamental aspect of our bodily self-consciousness (Blanke et al., 2004; Dijikerman, 2015), but also because the representation of one’s own body is not fixed but rather the result of a predictive processing that is constantly updated by means of multisensory processes. Future studies should explore and extend these findings to clinical populations characterised by body image distortions and lack of bodily awareness, such as eating disorders.

## Footnotes

**Authors Contributions** LC, AF and PMJ conceived and designed the experiment. LR offered medical supervision to the study. LC performed data collection and analysed the data, under supervision of AF, PJ and YP. LC, YP, AF and PMJ wrote the manuscript. All the authors approved the manuscript before submission.

## Acknowledgements

Authors would like to thank Dora Tamari-Tutnjević, Mengbing Zhang, Amanda Hornsby, Mariana von Mohr for assistance with data collection. We are grateful to Miss Catherine Maze for her help with the customized rubber hands. The study was supported by a “European Research Council Starting Investigator Award” [ERC-2012-STG GA313755] to A.K.; a University of Hertfordshire studentship to L.C. and a Neuropsychoanalysis Foundation grant to L.C. Funding for the time of A.K. and L.C has been partially provided by the Fund for Psychoanalytic Research through the American Psychoanalytic Association.

